# Evidence of a cellulosic layer in Pandoravirus tegument and the mystery of the genetic support of its biosynthesis

**DOI:** 10.1101/793711

**Authors:** Djamal Brahim Belhaouari, Jean-Pierre Baudoin, Franck Gnankou, Fabrizio Di Pinto, Philippe Colson, Sarah Aherfi, Bernard La Scola

**Affiliations:** Aix-Marseille Univ., Institut de Recherche pour le Développement (IRD), Assistance Publique - Hôpitaux de Marseille (AP-HM), MEPHI, 27 boulevard Jean Moulin, 13005 Marseille, France; IHU Méditerranée Infection, 19-21 boulevard Jean Moulin, 13005 Marseille, France

**Keywords:** Giant virus, Pandoravirus, amoeba, *Acanthamoeba*, *Megavirales*, capsid, tegument cellulose

## Abstract

Pandoraviruses are giant viruses of amoebae with 1 μm-long virions. They have an ovoid morphology and are surrounded by a tegument-like structure lacking any capsid protein nor any gene encoding a capsid protein. In this work, we studied the ultrastructure of the tegument surrounding Pandoravirus massiliensis virions and noticed that this tegument is composed of a peripheral sugar layer, an electron-dense membrane, and a thick electron-dense layer consisting in several tubules arranged in a helicoidal structure resembling that of cellulose. Pandoravirus massiliensis particles were stained by Calcofluor white, a fluorescent dye of cellulose, and the enzymatic treatment of particles by cellulase showed the degradation of the viral tegument. We first hypothesized that the cellulose tegument could be synthesized by enzymes encoded by Pandoravirus. Bioinformatic analyses revealed in Pandoravirus massiliensis, a candidate gene encoding a putative cellulose synthase, with a homology with the BcsA domain, one of the catalytic subunits of the bacterial cellulose synthase, but with a low level of homology. This gene was transcribed during the replicative cycle of Pandoravirus massiliensis, but several arguments run counter to this hypothesis. Indeed, even if this gene is present in other Pandoraviruses, the one of the strain studied is the only one to have this BcsA domain and no other enzymes involved in the synthesis of cellulose could be detected, although we cannot rule out that such genes could have been undetected among the large proportion of Orfans of Pandoraviruses. As an alternative, we investigated whether Pandoravirus could divert the cellulose synthesis machinery of the amoeba to its own account. Indeed, contrary to what is observed in the case of infections with other giant viruses such as mimivirus, it appears that the transcription of the amoeba, at least for the cellulose synthase gene, continues throughout the growth phase of envelopes of Pandoravirus. Finally, we believe that this scenario is more plausible. If confirmed, it could be a unique mechanism in the virosphere.

## Introduction

Giant viruses of amoebae are a group of complex viruses phylogenetically closely relative to the Nucleo – Cytoplasmic Large DNA Viruses (NCLDVs), within a new proposed order named Megavirales (Colson et al., 2013). Since the discovery of the first giant virus of amoebae in 2003 (La Scola et al., 2003), dozens of new giant viruses were described, considerably expanding our knowledge about their diversity, structure, genomics and evolution.

In 2013, two new complex giant viruses, named pandoraviruses, were described (Philippe et al., 2013). They replicate in *Acanthamoeba castellanii* and compose a new phylogenetic group of giant viruses of amoebae related to phycodnaviruses. The first isolate, named Pandoravirus salinus, originated from a marine sediment layer of the Tunquen River in Chile. The second isolate, Pandoravirus dulcis, was isolated from the mud of a freshwater pond in Australia. Pandoraviruses harbor specific morphological and genetic features, including ovoid-shaped particles with an ostiole-like apex and measuring ~1.0 μm in length and ~0.5μm in diameter, classifying them as the second largest particles after those of Pithovirus sibericum (Legendre et al., 2014). These viruses have a double-stranded DNA genome, up to 2.5 Mb (for Pandoravirus salinus), the largest genome ever described to date in the virosphere (Philippe et al., 2013). Subsequently, the nature of an endocytobiont of *Acanthamoeba* isolated few years before in Germany from the contact lens and storage case fluid of a patient with keratitis, was recognized as the third pandoravirus and was named Pandoravirus inopinatum (Scheid, 2016; Scheid et al., 2014). In 2015-2016, we isolated three new pandoravirus strains from sewage and soda lake water samples collected in Brazil, by co-culture on *A. castellanii*. These viruses were named respectively Pandoravirus massiliensis, Pandoravirus pampulha, and Pandoravirus braziliensis (Aherfi et al., 2018). Other recent prospecting studies reported the isolation of seven additional strains: Pandoravirus quercus, isolated from a soil sample collected in Marseille (France); Pandoravirus neocaledonia, isolated from the brackish water of a mangrove near Noumea airport (New Caledonia), Pandoravirus macleodensis, isolated from a freshwater pond near Melbourne (Australia) (Legendre et al., 2018), Pandoravirus celtis, isolated from a soil sample collected in Marseille (Legendre et al., 2019) and 3 new pandoraviruses isolated from water samples in Brazil (Pereira Andrade et al., 2019).

Although other giant viruses with a similar ovoid morphology have been described, including pithoviruses, cedratviruses or Orpheovirus (Andreani et al., 2016, 2018; Legendre et al., 2014), pandoraviruses have the intriguing particularity to harbor no hints of a known capsid protein and, at the same time, the absence in virions of any structure similar to that of a known capsid (Aherfi et al., 2018; Legendre et al., 2018; Philippe et al., 2013; Scheid, 2016). In addition, no capsid-resembling protein was identified by proteomics in the Pandoravirus salinus and Pandoravirus massiliensis virions (Aherfi et al., 2018; Philippe et al., 2013). Pandoravirus virions are wrapped in a ≈70-nm-thick tegument-like envelope, composed of three layers: one ≈20 nm-thick of light density; one intermediate of ≈25 nm-thick that appears darker and composed of fibrils; and an external layer ≈25 nm-thick with a medium density (Philippe et al., 2013). At one of the particle apex, an aperture of ¾70 nm in diameter opens the viral tegument. An internal lipid membrane is present beneath the tegument, thus delimiting the particle core. The tegument and the internal content of pandoravirus virions are synthesized simultaneously, from the aperture-harboring apex, during neo-virion synthesis in the cytoplasm of the amoebae. However, the nature of this tegument has remained unknown to date. Its deciphering is needed to refine the definition of these giant viruses.

In the present study, we aimed to characterize the nature of the tegument of pandoraviruses.

## Material and methods

### Transmission electron microscopy, electron tomography and scanning electron microscopy

For negative staining, a drop of purified Pandoravirus massiliensis particles fixed with 2.5% glutaraldehyde in 0.1M cacodylate buffer was adsorbed onto formvar carbon films on 400 mesh nickel grids (FCF400-Ni, EMS). Grids were stained for 10 seconds with 1% molybdate solution in filtered water at room temperature. For sections, pandoravirus particles were ultracentrifugated at 8,000 g for 10 minutes and fixed for 1h at 4°C under gentle mixing after pellet resuspension in a mixture of 1.2 % glutaraldehyde / 0.05% ruthenium red in 0.1M cacodylate buffer. Virus particles were ultracentrifugated at 8,000 g for 10 minutes to discard supernatant and were then fixed for 3h at 4°C in a mixture of 1.2 % glutaraldehyde / 0.05% ruthenium red in 0.1M cacodylate buffer. Then, Pandoravirus massiliensis virions were washed thrice 10 min with 0.1M Cacodylate buffer at 4°C. Next steps were performed at room temperature. Viral particles were rinsed twice for 15 min each, with a cacodylate 0.1 M / saccharose 0.2 M in water solution, and were dehydrated with ethanol 50%, 70% and 96%, for 15, 30 and 30 min, respectively. Pandoravirus massiliensis virions were then placed for 1h in a mix of LR-White resin 100% (Ref. 17411, MUNC-500; Polysciences) and ethanol 96% in a 2:1 ratio. After 30 min in pure 100% LR-White resin, particles were placed in 100% LR-White resin overnight at room temperature. Pandoravirus particles were then placed for 1h in 100% resin at room temperature. A total of 1.5 mL of Pure 100% LR-White resin was added on the virus pellet. Polymerization was achieved at 60°C for 3 days. Between all steps of inclusion, the samples were ultracentrifuged at 2,400 g and the supernatant was discarded. Sections of 70 or 300 nm-thick were cut on a UC7 ultramicrotome (Leica). Ultrathin sections were deposited on 300 mesh copper/rhodium grids (Maxtaform HR25, TAAB). They were post-stained with 5% uranyl acetate and lead citrate according to the Reynolds method (Reynolds, 1963). Gold nanoparticles with a diameter of 10 nm (Ref.752584; Sigma-Aldrich) were deposited on both faces on the 300 nm thick ultrathin sections for tomographic fiducial alignment. Electron micrographs were acquired on a Tecnai G^2^ transmission electron microscope operated at 200 keV and equipped with a 4096 × 4096 pixel resolution Eagle camera (FEI). Tomography tilt series were acquired with the Explore 3D (FEI) software for tilt ranges of 110° with 1° increments. The mean applied defocus was − 4 μm. The magnification ranged between 9,600 and 29,000 with pixel sizes between 1.12 and 0.37 nm, respectively. The average thickness of the tomograms obtained was 155 ±43 nm (n= 8 measures). The tilt-series were aligned using ETomo from the IMOD software package (University of Colorado, USA) (Kremer et al., 1996) by cross-correlation. The tomograms were reconstructed using the weighted-back projection algorithm in ETomo from IMOD (Kremer et al., 1996). ImageJ software was used for image processing (Schneider et al., 2012).

### Cellulose staining and light microscopy

Calcofluor staining was achieved by depositing a drop of purified pandoravirus particles in PAS medium onto a glass slide and immediately adding 50μL of calcofluor white (Ref.18909; Sigma-Aldrich) and 50 μL of 10% KOH prior to glass slide covering and confocal imaging. Pandoravirus particles were imaged with a Plan-Aprochromat x63/1.4 immersion objective on a AiryScan LSM800 confocal laser scanning microscope (Zeiss). Image size was 512×512 pixels and scan zoom ranged from x0.5 to x2.9. Laser excitation with a 405 nm wavelength was used for calcofluor staining imaging and was coupled to an ESID detector for depicting particles contours.

### Enzymatic treatment of Pandoravirus massiliensis virions

A total of 50 μL of purified virions was added to 1 mL of cellulase solution (cellulase from *Trichoderma reesei*, aqueos, Sigma Aldrich, C2730-50ML) at different concentrations and incubated for 48 h at 45°C. In a second time, these samples were imaged on AiryScan LSM800 microscope confocal laser scanning microscope. The effects were then observed by confocal microscopy (AiryScan LSM800), scanning microscopy and transmission electronic microscopy. The number of intact viral particles was estimated using the imageJ software (Schneider et al., 2012)

### Bioinformatic analyses to search for cellulose synthase candidate genes in the Pandoravirus massiliensis genome

Sequences of the predicted ORFs of Pandoravirus massiliensis in amino acids were used for BLAST searches against the NCBI GenBank nr database. The analyses were performed using 1e-2, 25% and 50% as thresholds for the evalue, the homology and the coverage of aligned sequences, respectively. Phylogenetic reconstruction was performed using the Maximum Likelihood method with the MEGA6 software (Tamura et al., 2013). Conserved domains were also searched for by DELTA-BLAST analyses against the Conserved Domain Database (Marchler-Bauer et al., 2017).

BLASTp, tBLASTn and BLASTn analyses were also performed against the gene contents and complete genomes of the 9 other pandoraviruses, i.e. Pandoravirus dulcis, Pandoravirus salinus (Philippe et al., 2013), Pandoravirus inopinatum (Scheid, 2016; Scheid et al., 2014), Pandoravirus quercus (Legendre et al., 2018), Pandoravirus macleodensis (Legendre et al., 2018), Pandoravirus neocaledonia (Legendre et al., 2018), Pandoravirus celtis (Legendre et al., 2019), Pandoravirus braziliensis and Pandoravirus pampulha (Aherfi et al., 2018).

### Transcription of candidate gene of cellulose synthase of Pandoravirus massiliensis and cellulose synthase of *Acanthamoeba castellanii*

RNA was extracted with the RNeasy mini kit (Cat No: 74104, Qiagen, France) at different time points of the Pandoravirus massiliensis cycle, from H0 (i.e. 45 min after the inoculation of amoeba cells by viral particles) until H12 post-infection (release of neo-synthetized virions). Total RNA was eluted in 50 μL of RNase-free water; 2.5 μl of RNaseOUT (Thermo Fisher Scientific, France) was added to the eluate to discard RNase. The DNase treatment was performed with TURBO DNase (Invitrogen, France; six cycles of 30 min incubation at 35°C). Two PCR systems targeting the DNA polymerase gene of Pandoravirus massiliensis (forward primer: 5’-ATGGCGCCCGTCTGGAAG; reverse primer: 5’-GGCGCCAAAGTGGTGCGA) and the housekeeping gene of the RNA polymerase of Acanthamoeba castellanii (forward: 5’-ACGAACTTCCGAGAGATGCA; reverse: 5’-CACCTTGACCAGTCCCTTCT) were used to check for genomic DNA contamination and as positive controls for the reverse transcription. Primers targeting the candidate gene for the putative cellulose synthase of Pandoravirus massiliensis (forward: 5’ TCCACTCGACATGCAATCTT; reverse: 5’-AAAACACAAACCCGCTCTGC) and those targeting the cellulose synthase genes (KC466026.1 and XM_004335119.1) of Acanthamoeba castellanii (forward:5’GGGAGATCAACGACAACCTG; reverse: 5’-GTCCTCRGTCTGCGACTCGT) were designed using Primer3 (Koressaar and Remm, 2007). RNeasy MinElute Cleanup Kit (Qiagen) was used to purify total RNA according to the manufacturer’s recommendations. Total RNA was reverse-transcribed into cDNA by using the SuperScript VILO Synthesis Kit (Invitrogen, France). Then, qPCR was carried out on the cDNA with the LightCycler^®^ 480 SYBR Green 1 Master reaction mix (Roche Diagnostics, Mannheim, Germany), following the manufacturer’s temperature program with 62°C as primer hybridization and elongation temperature. Each experiment was performed in triplicate.

## Results

### Pandoravirus massiliensis ultrastructure

The study by transmission electron microscopy (TEM) of the ultrastructure of negatively stained purified Pandoravirus massiliensis particles showed ovoid shaped virions with a mean maximal diameter of 1,230±179 nm and a mean minimal diameter of 689±114 nm (n=10), these dimensions reaching up to 1,510 nm × 860 nm (Figure 1.A). An ostiole with a concave shape could be observed at one apex of the particles (Figure 1.A) and thin fibrillar structures were present around the particles. Fixation with ruthenium red allowed good visualization of peripheral polysaccharides on ultrathin sections. We observed for all particles, from periphery to inside (Figure 1.B,C): (i) peripheral sugars as depicted by ruthenium red aggregates, with electron-dense spikes originating from a thick layer of electron-dense aggregates; (ii) an electron-lucent space; (iii) an electron-dense membrane; (iv) an homogeneous interspace; (v) a thick electron-dense layer made of several tubules; (vi) a smooth internal compartment, more dense at each apex. Particles were cut along different planes, thus showing their different orientations, and the ostioles could be observed cut transversally or perpendicularly (Figure 1.D). Structures originating from the thick inner tubular layer could be observed, such as thick tubules reaching the thin outer electron-dense membrane (Figure 1.E) or thin fibrils reaching the most peripheral ruthenium-red stained polysaccharides (Figure 1F).

**Figure 1.**
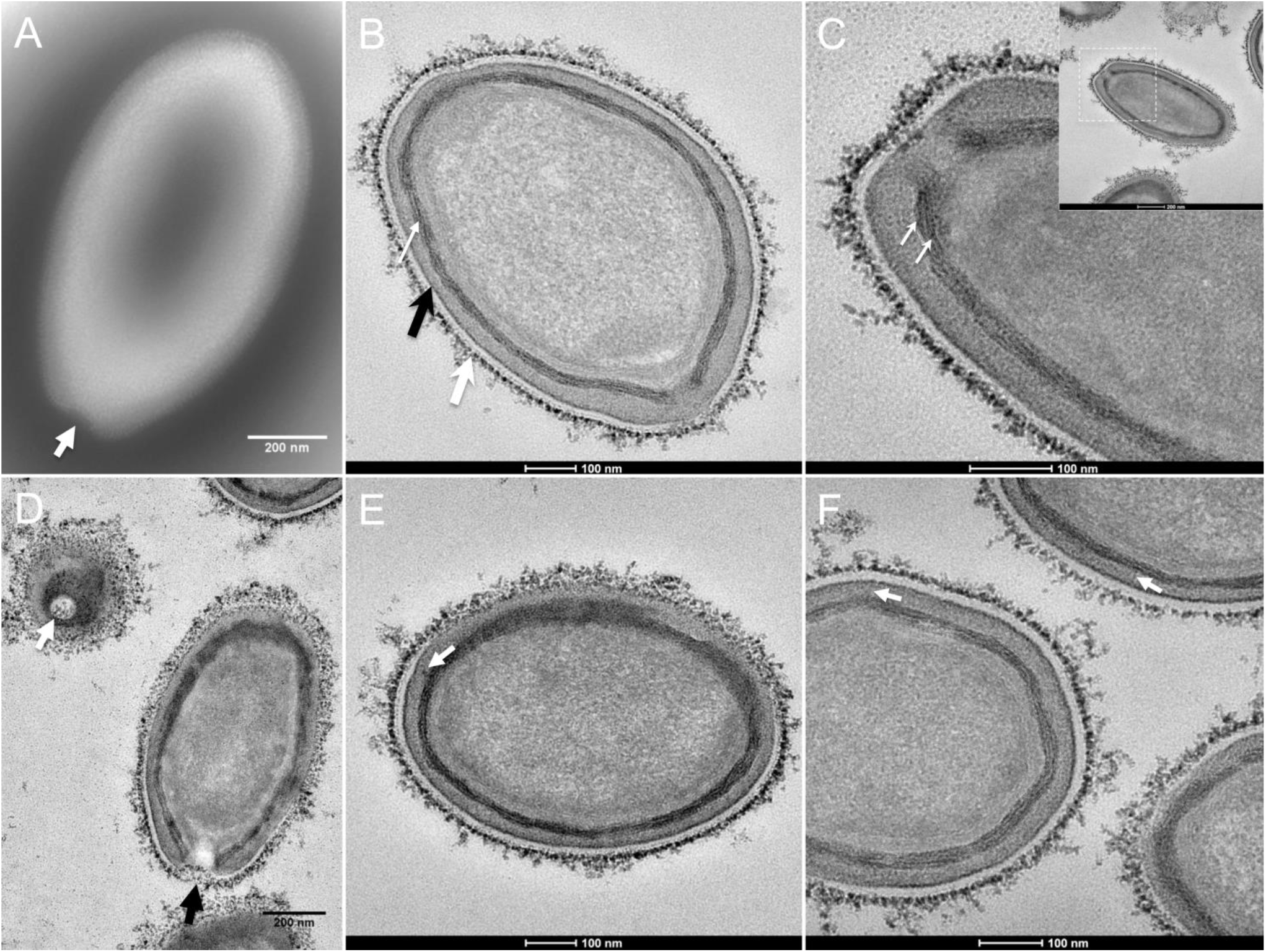
Transmission electron microscopy of Pandoravirus massiliensis. A. Negative staining of a Pandoravirus massiliensis particle: the ostiole (arrow) is located at the apex of the particle. Peripheral thin fibers can be observed enwrapping the particle. B. Ultrathin section showing i) the most-peripheral sugars depicted by ruthenium red aggregates (thick white arrow); ii) a thin electron-dense membrane (thick black arrow) and more centrally iii) a thick bundle of tubules (thin white arrow). C. Two thick tubules compose the inner-most thick layer. D. Two particles with ostioles cut transversally or perpendicularly. E. The inner-most thick tubules with thick protrusions toward the outer thin electron-dense membrane. F. Thin fibers projecting from the inner-most thick tubules, crossing the outer electron-dense membrane and reaching the peripheral ruthenium-red stained sugars

Next, to get a more detailed ultrastructure, a three-dimensional (3D) electron tomography on 300 nm-thin sections of ruthenium-red fixed LR-White embedded Pandoravirus massiliensis particles was performed. Eight tomograms were reconstructed. A tilt-series acquired at x14,500 magnification (Movie 1) was used to reconstruct Tomogram 1, in which several particles can be observed (Movie 2). These particles were ovoid in shape with a homogeneous internal compartment, or non-ovoid with an electron-luscent internal compartment, suggesting deteriorated particles. Tomogram 2 (Movie 3) is a zoom-in on a particle from Tomogram 1. Selected Z-planes from tomogram 2 (Figure 2) illustrate structures originating from the inner thick tubular layer: thick tubular structures with various diameters (black arrows in Figure 2) and thinner tubular/membranous structures of 2 nm-thick at the the ostiole (white arrows in Figure 2). Tomogram 3 is shown in Movie 4. Selected Z-planes from tomogram 3 in Figure 3 show that the inner-most thick tubular layer is composed of two tubules with a diameter of 8 nm (Figure 3.B). Thin 2 nm-thick fibrils originate from the tubular layer, crossing the electron-dense outer membrane and projecting toward the peripheral sugars (Figure 3.C), or oriented toward the internal compartment on the opposite side of the ostiole (Figure 3D). Tomogram 4 (Movie 5 and Movie 6) was chosen to illustrate the continuity between the two tubules from the inner-most tubular layer. These tubules can form a U-shape with no ending at the ostiole (Figure 4.A). In addition, we found out that the two inner-most tubules can present a helicoidal arrangement. This finding is shown in tomogram 5 (Movie 6), which is a zoom-in sub-tomogram from tomogram 4. In Figure 4.B, the two tubules from tomogram 5 are arranged as a helix, with locations where the two tubules are distant and other locations where the two tubules are crossed. This helical structural arrangement of the inner-most thick tubular layer then became obvious when playing all tomograms acquired, and this organization could also be noticed when looking back at conventional ultra-thin sections without tomography. The two tubules were superposed forming a 10 nm-thick layer or alternatively distant of 30 nm. On average, this helical arrangement had a periodicity of 150 nm, from crossing points to the most distant location point of the tubules.

**Figure 2.**
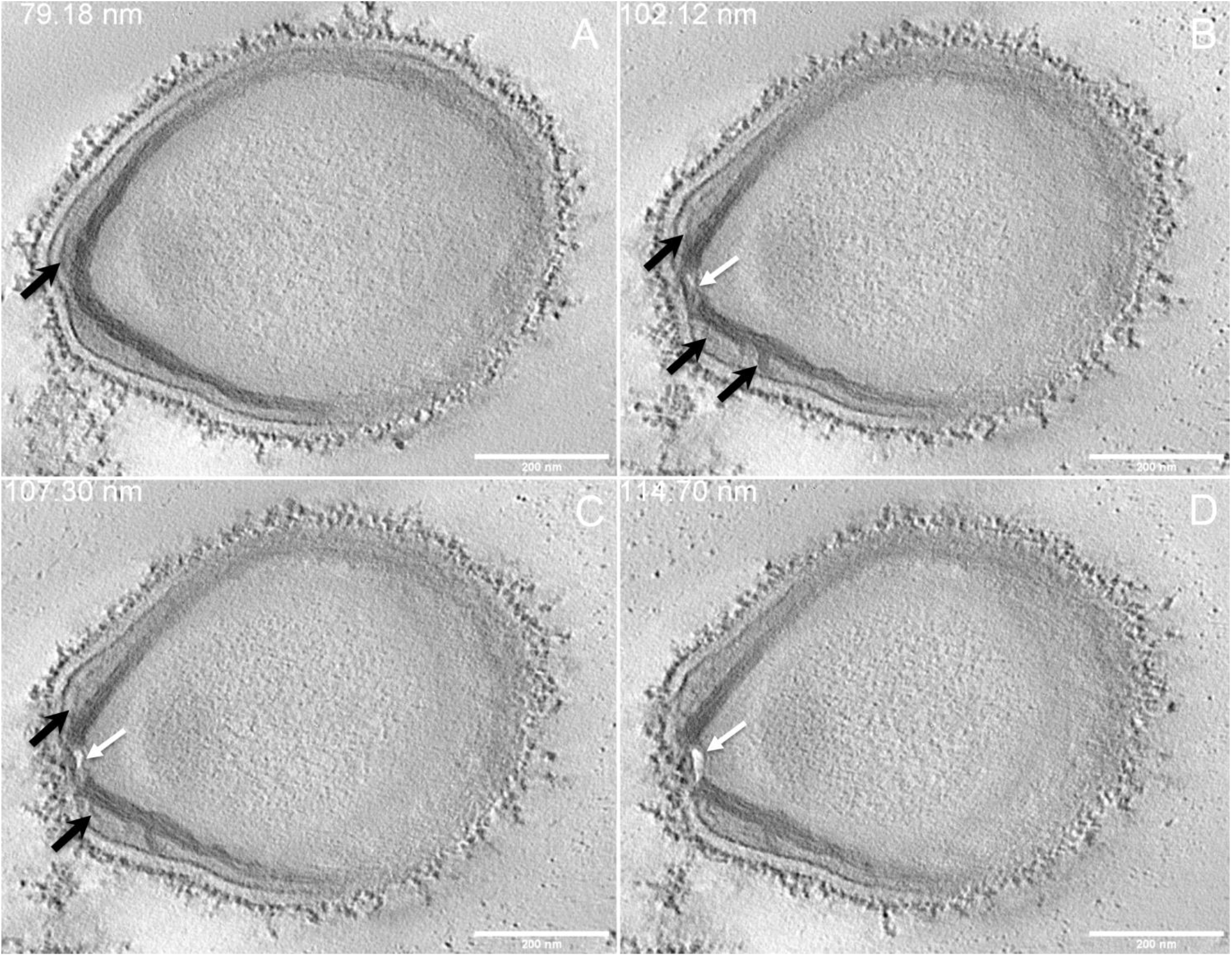
Electron tomography of Pandoravirus massiliensis particle from Movie3. A-D. Single planes in the tomogram from Movie3 showing thick tubules protruding toward the periphery and the outer electron-dense membrane (black arrows); a thin tubule/membrane (white arrow) connects the thick tubules layers located on each side of the ostiole.

**Figure 3.**
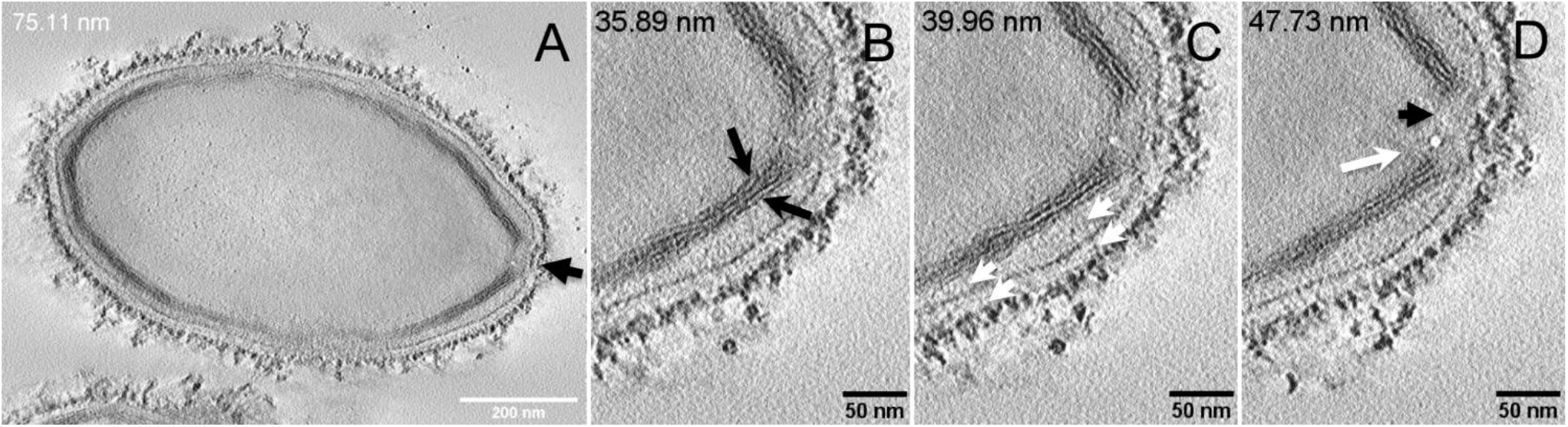
Electron tomography of Pandoravirus massiliensis particle from Movie4. A. Single plane in the tomogram from Movie4 showing a whole Pandoravirus particle and its ostiole located at one apex (arrow). B. The inner-most layer is composed of two thick tubules well separated as seen here or contacting each other. C-D. Thin fibers (white arrows) originating from the inner-most tubular thick layer projecting toward the peripheral sugars (white arrows, C), toward the inner core of the particle (white arrow, D) or at the level of the ostiole (black arrow, D).

**Figure 4.**
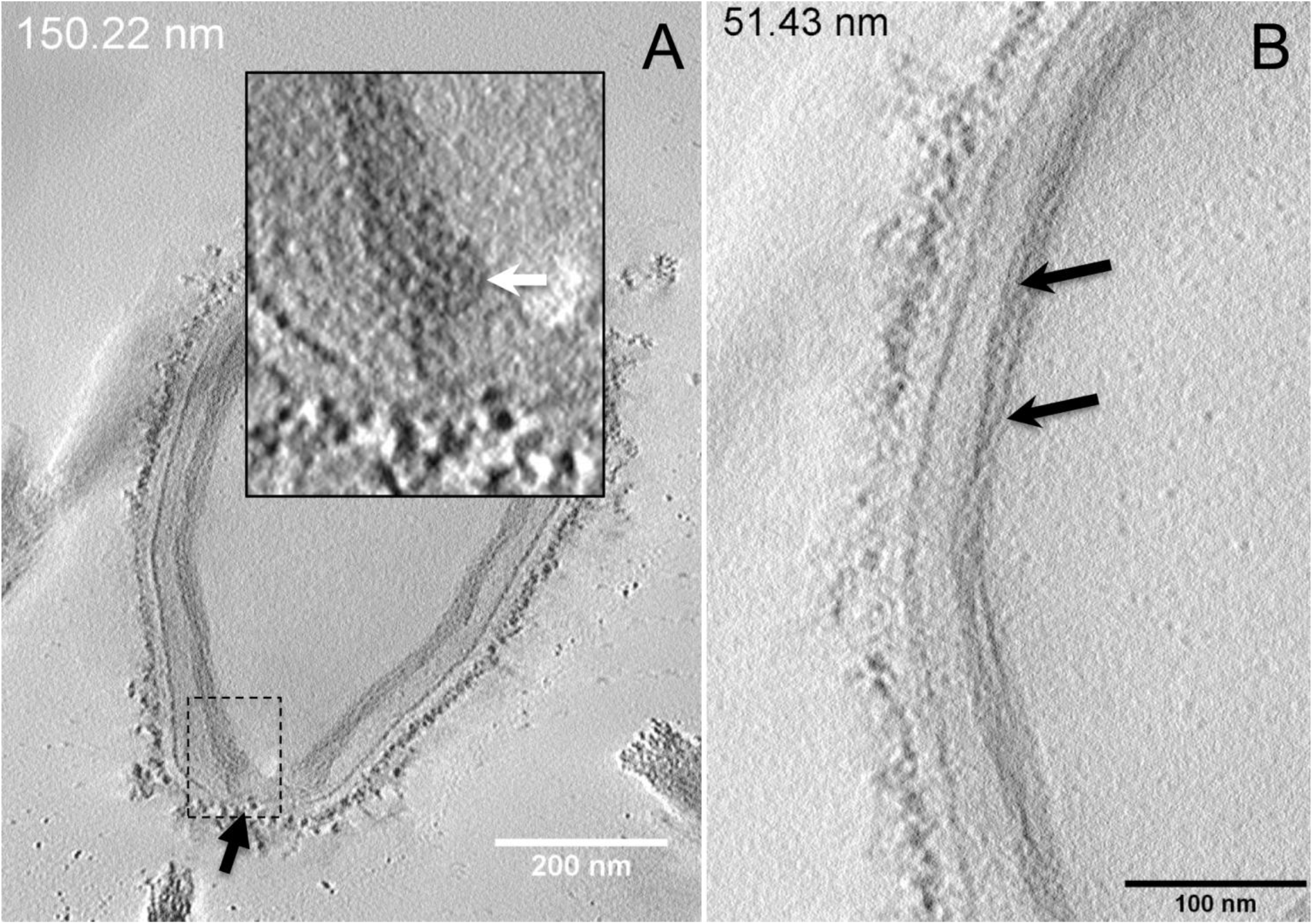
Electron tomography of Pandoravirus massiliensis particles from Movies 5 and 6. A. Single plane in the tomogram from Movie5 showing a Pandoravirus particle with its ostiole (black arrow). The magnified boxed region depicts a U-shaped thick tubule from the inner-most layer. B. Single plane in Movie6 from the zoomed-in tomogram from Movie5 showing the helical structural arrangement of the two thick tubules (black arrows) composing the inner-most layer of Pandoravirus particles.

Subsequently, since the diameter and structure of inner-most tubules potentially forming helix resembled cellulose and chitin according to the literature, we hypothesized thatthat the inner-most layer of the Pandoravirus massiliensis particles was consisted of cellulose and/or chitin.

### Cellulose staining

In order to check for cellulose content in Pandoravirus massiliensis particles, a calcofluor white staining of viral particles smears and confocal imaging were performed. Negative control consisting in unstained particles did not show any fluorescence under UV illumination (Figure 5.A, B). Inversely, calcofluor-stained particles were fluorescent under the same illumination (Figure 5.C, D). Single in-liquid Pandoravirus massiliensis particles were stained by calcofluor, as well as particles that remained in a few amoebae still present after purification (Figure 6.A). Zooming on individual particles showed that, on average, the fluorescence of the periphery of the particles seemed to be more intense than in the central region of the same particles (Figure 6.B, C).

**Figure 5.**
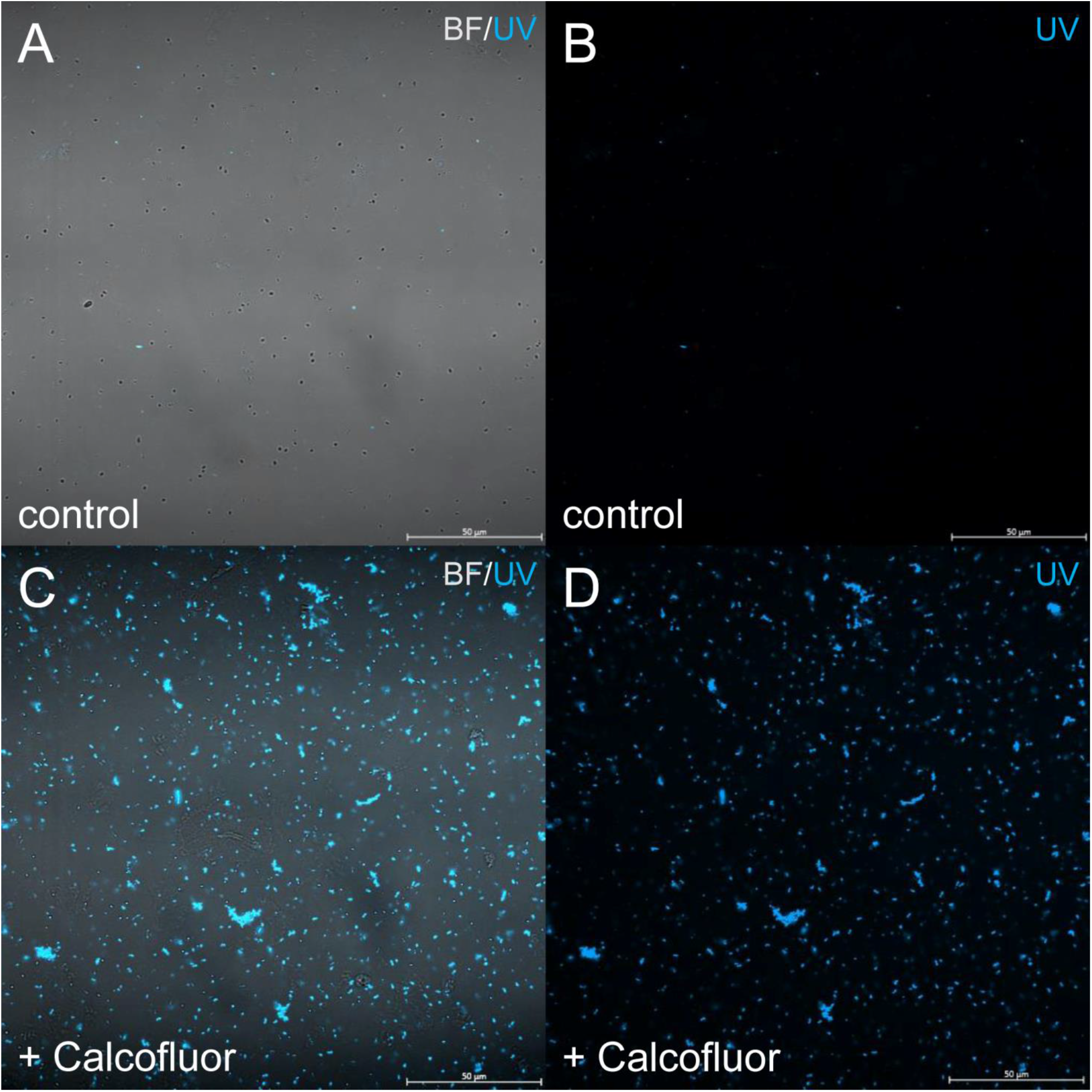
Confocal imaging of Calcofluor staining of Pandoravirus massiliensis. Control Pandoravirus particles. C,D. Calcofluor-stained Pandoravirus particles.

**Figure 6.**
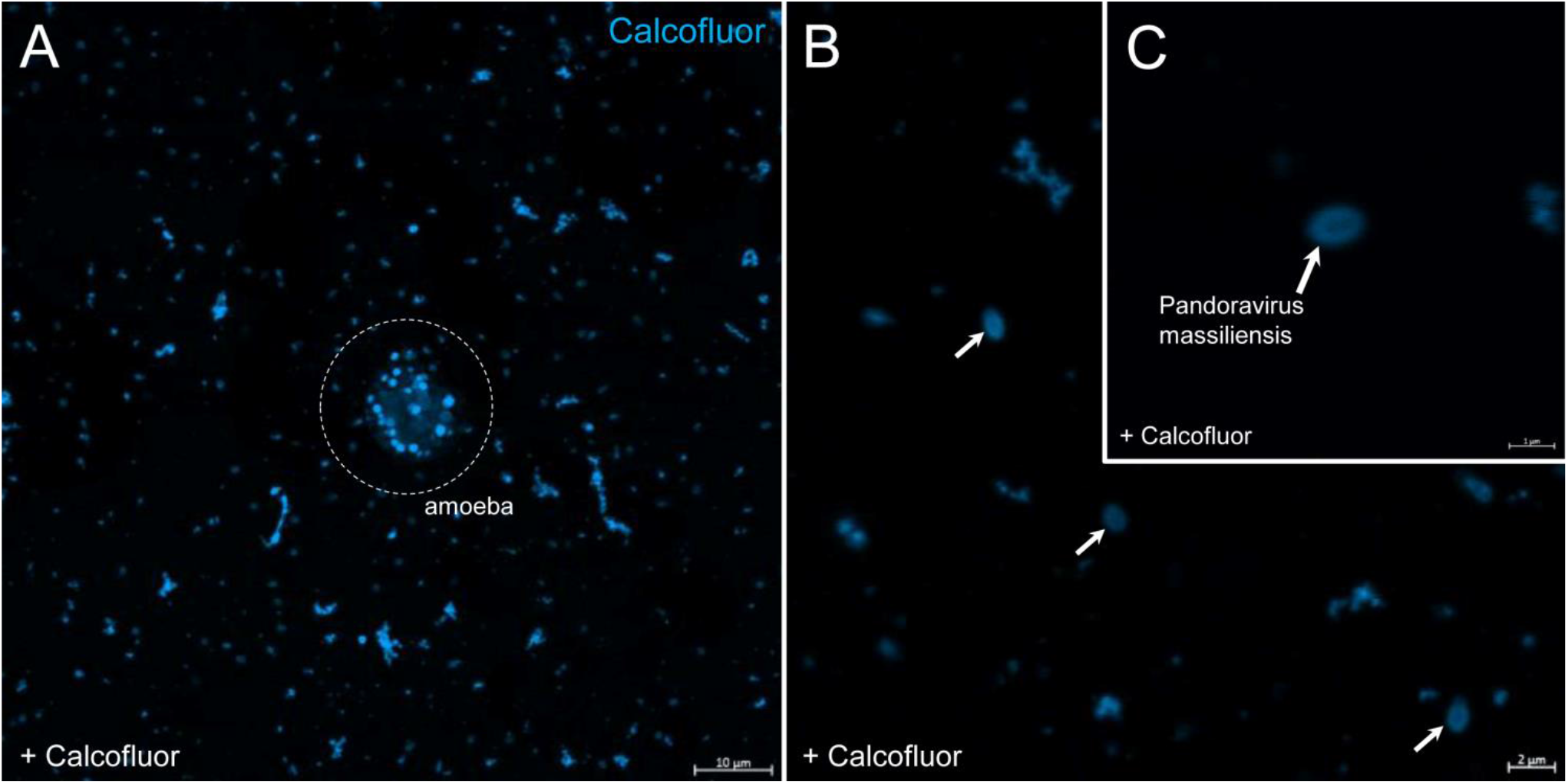
Confocal imaging of Calcofluor staining of Pandoravirus massiliensis. A. Pandoravirus-infected amoeba and single Pandoravirus particles stained with Calcofluor white. B, C. Calcofluor-stained Pandoravirus particles showing an intense peripheral calcofluor signal and a less-stained central region.

### Degradation of the pandoravirus particles by the cellulase treatments and ultrastructure of cellulase-treated Pandoravirus massiliensis particles

The mean number of viral particles per microscopic field observed after cellulase treatment at high concentrations (stock solution and 1:10 dilution) by confoncal microscopy and assessed on 10 microscopic fields by the ImageJ software decreased in comparison with the negative control composed of the same viral sample not submitted to the enzymatic treatment (Figures 7, 8). With a cellulase solution diluted at 1:10, the number of viral particles was slightly divided by two, and only ≈6% of the virions remained intact after a treatment by the cellulase stock solution. Moreover, calcofluor staining on viral particles treated by cellulase did not show any fluorescence under UV illumination (Figure 9. A1). Besides, a treatment by 1:10 diluted cellulase solution showed a fluorescent matrix (Figure 9.B1) suggesting a degradation of a cellulosic part of the viral particles. It should be noted that this result was less obvious in the most diluted solution of the enzyme (1:100) (Figure 9.C1). Furthermore, the scanning microscopy of pandoravirus virions treated with cellulase at high concentration confirmed the progressive degradation of particles (Figures 9.A1 to D3). It also showed the partial digestion of pandoravirus particles treated with cellulase at 1:10 dilution, by forming a matrix (Figures 9.B2,B3).

**Figure 7.**
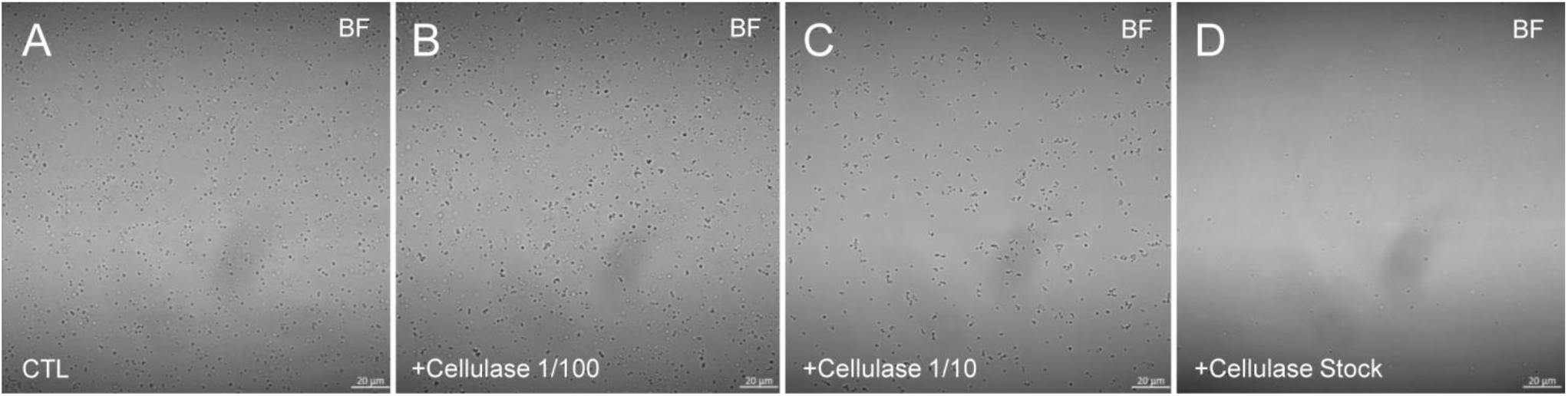
Confocal imaging of cellulase-treated Pandoravirus massiliensis. A. Control condition with untreated Pandoravirus massiliensis particles. B-E: cellulase-treated Pandoravirus massiliensis particles.

**Figure 8:**
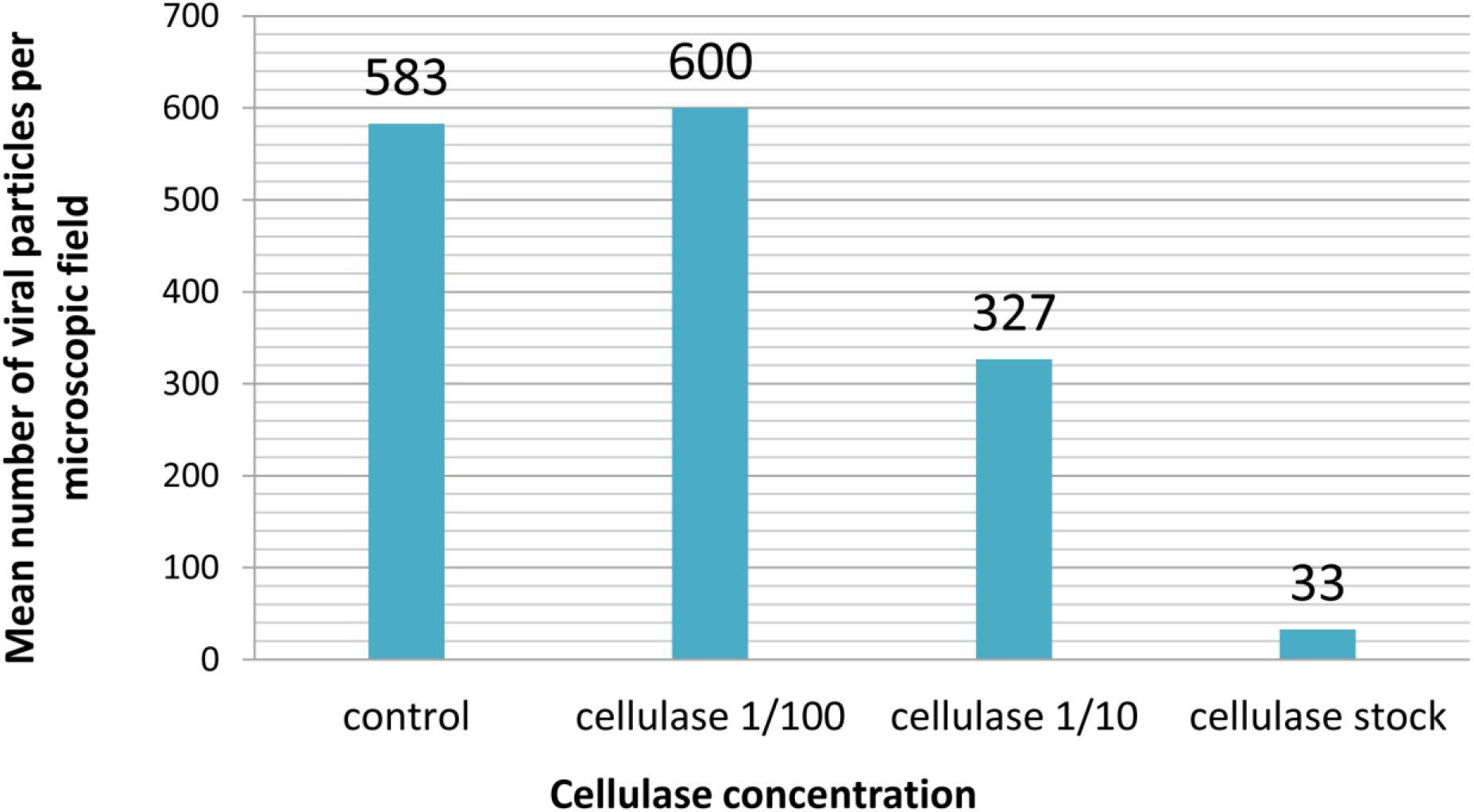
Estimation of the mean number of particles of Pandoravirus massiliensis per microscopic field of observation after cellulase treatment, assessed by the imageJ software.

**Figure 9.**
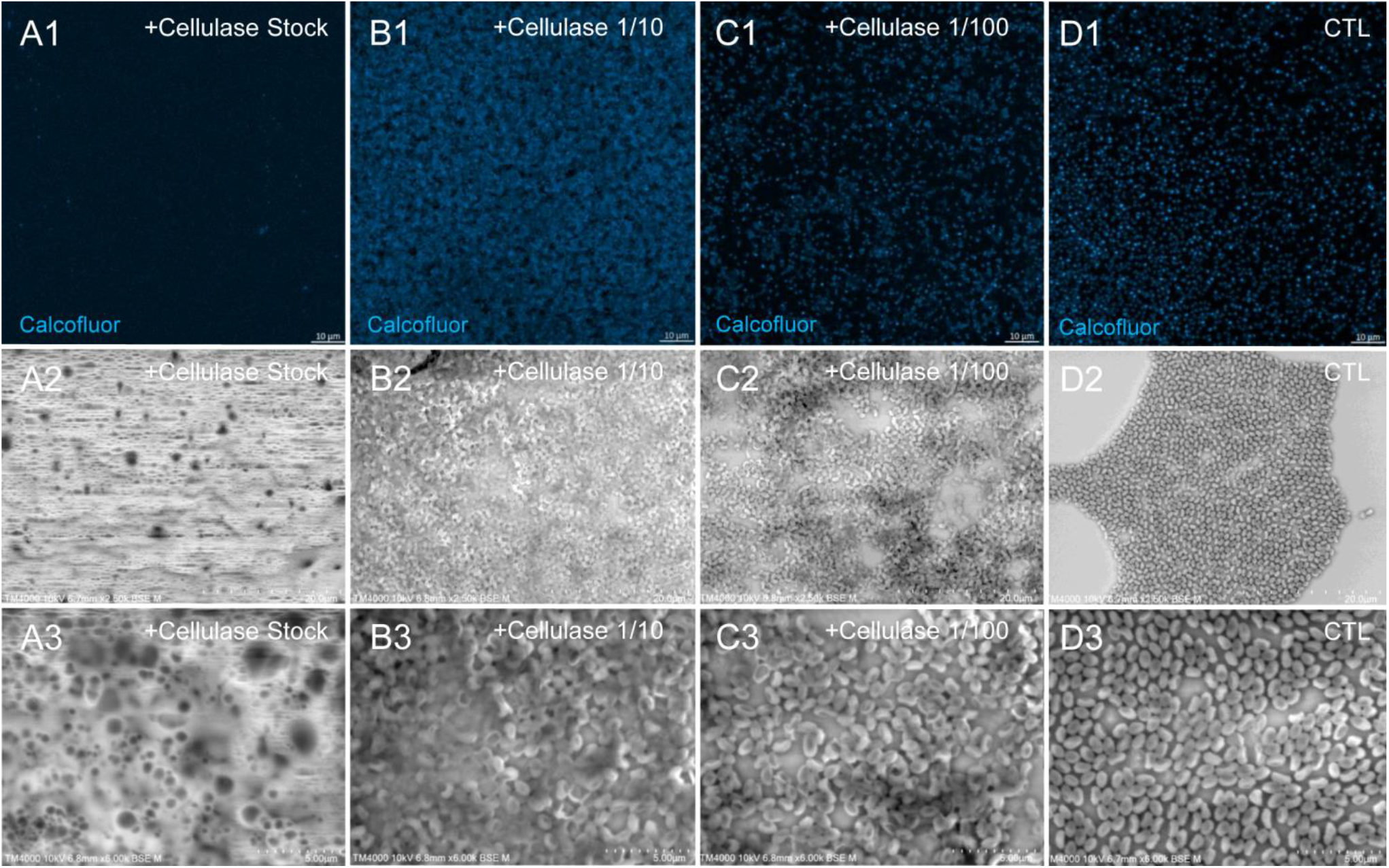
Confocal imaging of Calcofluor-stained cellulase-treated Pandoravirus massiliensis virions and scanning microscopy of cellulose treated Pandoravirus massiliensis. A1-C1: cellulase-treated Pandoravirus massiliensis particles stained with Calcofluor-white.D1 Control condition with untreated Pandoravirus massiliensis particles stained with Calcofluor-white. A(2-3) B(2-3)-C(2-3): cellulase-treated Pandoravirus massiliensis particles.D2.D3 Control condition with untreated Pandoravirus massiliensis .. imaged with scanning microscopy on two magnification.

As compared to control, untreated Pandoravirus massiliensis particles (Figure 10.A1-A5), a degradation of the virions by the cellulase solution was observed by transmission electron microscopy (Figure 10.B-D). Albeit this effect was not homogeneous between particles in all conditions, defective particles were observed with a dose-dependent effect of cellulase at the peripheral envelope and/or the internal compartment. Indeed, at 1/100 cellulase concentration, particles showed a defect of the envelope (Figure 10.B3-B5) with the presence of electron-luscent regions between the thin electron-dense membrane and the thick bundle of tubules described in Figure 1. Detachments from the different layers were also observed in most of the affected particles.

**Figure 10.**
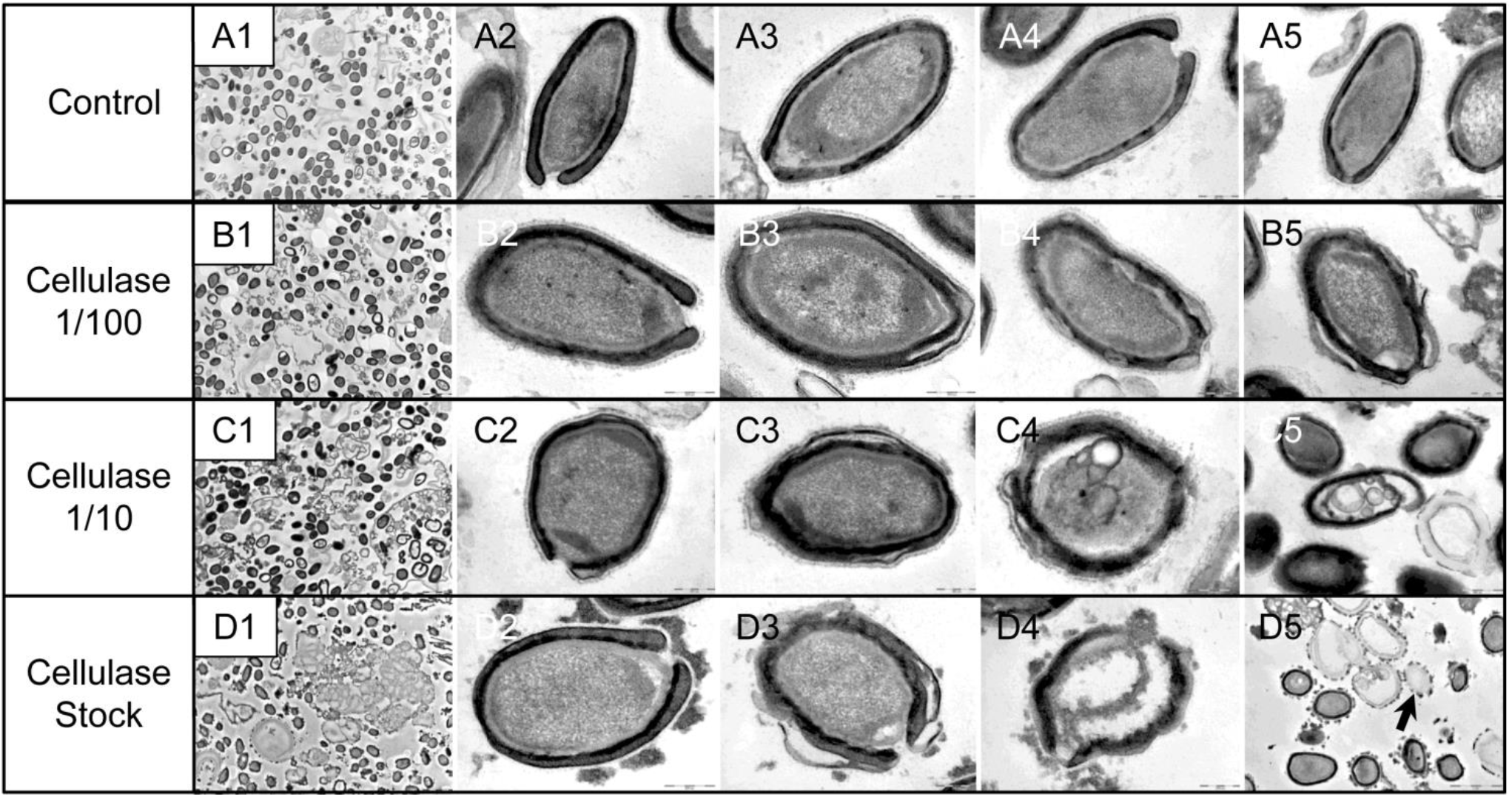
Scanning microscopy of cellulase-treated Pandoravirus massiliensis particles. A1-A4: Control condition with untreated Pandoravirus massiliensis particles. B-D Cellulase-treated Pandoravirus massiliensis particles imaged in different stages of digestion 1-4 from least to the most digested B.C.D (1): intact particles. B.C.D (2): the beginning of digestion. B.C.D(3): in the middle of digestion. B.C.D (4): at the end of the enzymatic digestion.

(Figure 10.B5). Accordingly, with 1/10 cellulase concentration treatment, particles exhibited electron-luscent regions at the level of the envelope as well as detachments of the different layers (Figure 10.C3-C5), and also defects in the internal compartment with empty spaces and vacuoles (Figure 10.C4,C5). This defect in the internal compartment was even more observable at stock concentration of cellulase (Figure 10.D2-D4), with ‘ghost-like’ particles presenting totally empty internal spaces and only a thin surrounding envelope (Figure 10.D5). We also noticed the presence of debris and/or of an amorphous matrix in the ultra-thin sections of cellulase-treated pandoravirus virions (Figure 10.B1,C1,D1), in a dose-dependent manner coherent with the amorphous matrix seen by light microscopy and scanning electron microscopy under the same conditions (Figure 10), which may correspond to the debris of cellulase-digested particles.

### Cellulose synthase candidate gene in the Pandoravirus massiliensis gene content

DELTA-BLAST analyses revealed a distant homology for the predicted gene 594 of Pandoravirus massiliensis with the cellulose synthase domain bcsA of different bacteria such as *Escherichia coli*, *Shigella* sp., *Salmonella enterica* for the 30 best hits. These homologies were barely significant with the 10 best hits, identity percentages in amino acids varying between 24.7% and 23.9%, query coverages between 24% and 25%, and e-values ranging between 2e-15 and 3e-15 (Supplementary File 1). For this ORF594, no homology with a cellulose synthase was found by BLAST analyses neither against the nr database, nor against the conserved domain database (CDD) (Marchler-Bauer et al., 2017). Orthologs of this ORF594 were found by BLASTp analyses in all the 9 other pandoraviruses, with e-values varying from 3.41e-11 to 0.006; but none of them harbored the same bcsA domain when searched for by DELTA-BLAST analysis or in the Conserved Domain Database.

### Transcription of candidate gene of cellulose synthase of Pandoravirus massiliensis and cellulose synthase of *Acanthamoeba castellanii*

All qPCR experiments performed on *P. massiliensis* DNA using primers targeting the cellulose synthase candidate gene (ORF594) were positive, whereas all those performed on *A. castellanii Neff* DNA using the same primers were negative, which confirmed the specificity of the PCR system. Conversely, qPCR targeting the cellulose synthase gene of *A. castellanii* was positive on the amoebal DNA and negative on the viral DNA, confirming the amoebal specificity of these primers. All qPCR tests performed on the purified RNA extract after DNase treatment before the reverse transcription step were negative, indicating efficient DNA degradation.

qRT-PCR targeting *P. massiliensis* ORF594 were positive on the viral cDNA for samples collected during the whole viral cycle, indicating that this gene was transcribed. qRT-PCR targeting *A. castellanii* cellulose synthase gene were positive on the cDNA obtained from uninfected amoebae and from amoebae infected by *P. massiliensis* during the whole replicative cycle (Figure 11). These latter results show that the cellulose synthase gene of *Acanthamoeba castellanii* is transcribed both in uninfected amoebae and in amoebae infected with Pandoravirus massiliensis.

**Figure 11.**
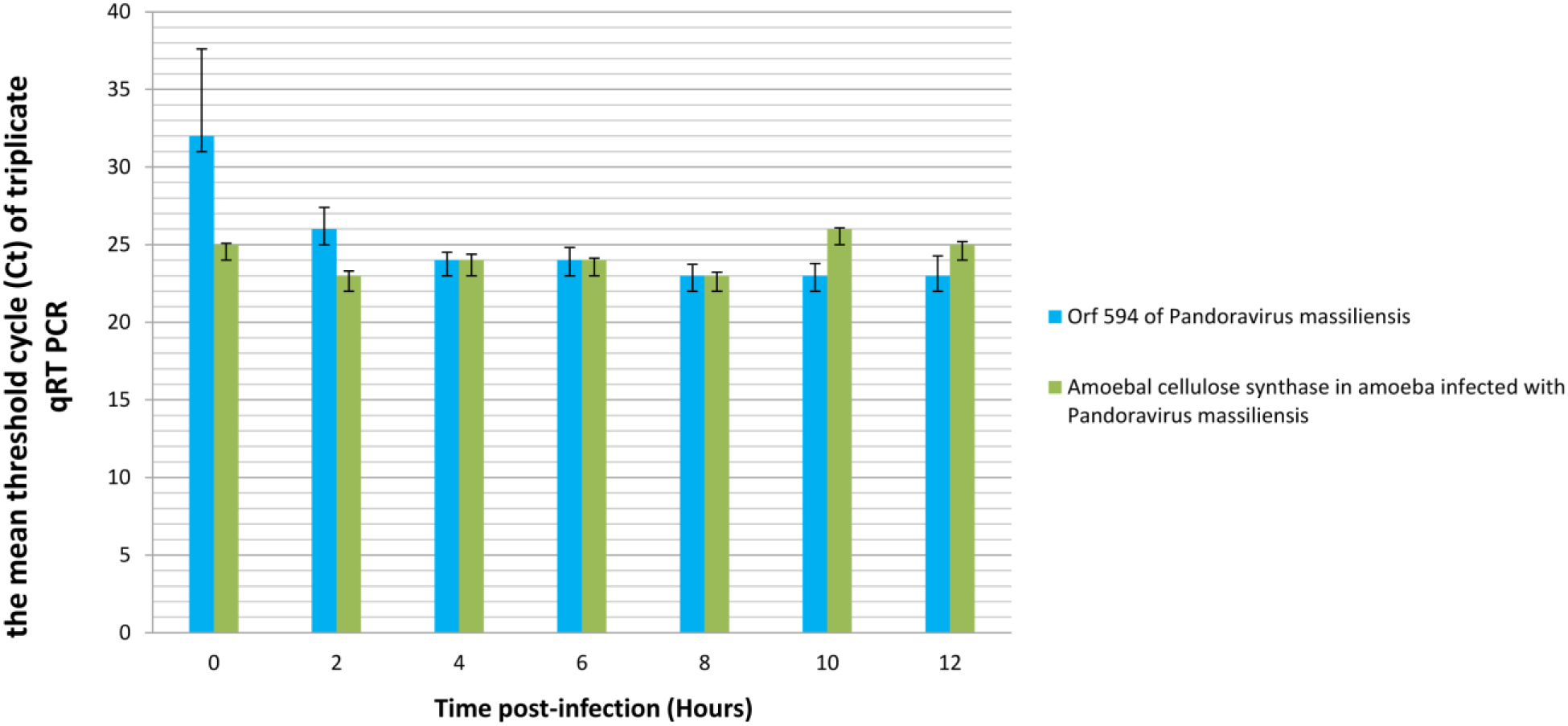
Representation of the mean threshold cycle (Ct) of triplicate qRT PCR on the RNA of Pandoravirus massiliensis by targeting the predicted gene of the cellulose synthase (ORF594) and on RNA of *Acanthamoeba castellanii* infected with Pandoravirus massiliensis by targeting the amoebal gene of the cellulose synthase, according to the time post-infection from 0 to 12 hours postinfection).

## Discussion

We have determined here, by studying the ultrastructure of Pandoravirus massiliensis virions by transmission electron microscopy after various treatments, that its viral integument, not previously characterized, was partially of a polysaccharide nature with a helical structure comparable to that of vegetable cellulose. Several markers of sugars (red rhutenium, calcofluor), the degradation of virions by cellulase and the use of appropriate negative controls clearly confirmed the cellulosic nature of the pandoravirus viral tegument. The cellulose is the most common biologic macromolecule, biosynthesized by vegetals, algae, some bacteria, and by some marine animals as Ascidiae (Chen et al., 2016; Helenius et al., 2006; Nakashima et al., 2004). It is also a major component of the cyst of Acanthamoeba, the unique host of pandoraviruses demonstrated to date (Magistrado-Coxen et al., 2019). It is composed of polymers of β (1,4) glucose subunits. Cellulosic chains are structured as microfibrils, which confer both resistance and plasticity to the vegetal walls, and also probably to the tegument of Pandoravirus massiliensis virions. The bioinformatic analyses of the Pandoravirus massiliensis genome possibly revealed a candidate gene (ORF594) for a cellulose synthase, which could be involved in the cellulose synthesis of the viral tegument. Indeed, a distant homology with the BcsA domain, which is one of the four catalytic subunits of the bacterial cellulose synthase was found (Wong et al., 1990). The domain bcsA polymerizes 5’-UDP-glucose to the cellulose polymer in formation (Zimmer, 2015). In the 9 other pandoraviruses, an ortholog for the ORF594 was found by BLASTp analysis. The experiments of qRT PCR revealed that this gene was transcribed during the viral cycle with an increased transcription 4, 6 and 8 hours post-infection, the time points matching with the formation of viral factories, which could reinforce this first hypothesis.

However, the similarity with the bcsA domain is very low and this domain was not found in none of the 9 other pandoraviruses. Besides, the length of this domain bcsA found in bacteria ranges approximately around 800 amino-acids in length. The ORF594 is slightly smaller with a predicted encoded protein of only 135 amino-acids in length. The 3 other subunits of the cellulose synthase were not found in the Pandoravirus massiliensis genome, neither by BLASTp nor by DELTA Blast analyses. Therefore, although this ORF594 is transcribed during the replicative cycle, the implication of the ORF594 in the synthesis of the cellulose is unlikely. One of the most amazing features of the pandoraviruses is the tremendous proportion of ORFans (genes without any homolog in the international sequence databases) and genes predicted to encode hypothetical proteins in their genome. These recently described viruses are so far to be exhaustively characterized. It has been shown that these hypothetical proteins are transcribed and translated (Aherfi et al., 2018; Legendre et al., 2018; Philippe et al., 2013; Scheid, 2016) suggesting that they have an efficient function for the virus. Among all these predicted genes, some of them could be implied in the synthesis of the cellulose, i.e. other unidentified cellulose synthase subunits, through synthesis by new enzymes or by a new metabolic pathway, unknown to date.

We could alternatively hypothesize that the virus could use for the synthesis of its particles tegument, the gene of the cellulose synthase of *Acanthamoeba castellanii* (Clarke et al., 2013). In a previous work, it has been shown that when *Acanthamoeba* was infected with a mimivirus, the transcription of the amoeba felt dramatically and at 6 hours after infection, the transcription became undetectable (Legendre et al., 2010). We observed herein, that the host gene of cellulose synthase was transcribed during the all the replication cycle of Pandoravirus massiliensis, showing that the transcription of this amoebal gene is not impacted by the infection with Pandoravirus. Therefore, we can assume that the amoebal cellulose synthase could be diverted to the virus benefits, and could be involved in the synthesis of the cellulosic viral tegument. This gene was also found as transcribed, while *Acanthamoeba castellanii* were in the survival buffer, a buffer with the minimal components for the survival of amoebae, and thus leading to the amoebal enkystment. At this time, the amoeba could start the synthesis of cellulose, a main component of the amoebal cyst.

In conclusion, we could determine herein the cellulosic nature of the tegument of Pandoravirus massiliensis. Although a distant similarity was found with the catalytic subunit bcsA of the bacterial cellulose synthase, with a predicted ORF of Pandoravirus massiliensis, this domain was not found in any other pandoravirus. These data suggest that the cellulose of the tegument of pandoraviruses could be probably the product of the host amoebal cellulose synthase. These results provide new data information about this important virus and contributes to the understanding of the biology of these complex viruses and their definition and classification in the virosphere.

## Authors contributions

DBB and JPB did the experiments and wrote the paper. FDP and FG did the experiments. PC wrote the paper. SA supervised the experiments and wrote the paper. BLS conceived the project, supervised the experiments and wrote the paper.

## Conflict of Interest

The authors declare that the research was conducted in the absence of any commercial or financial relationships that could be construed as a potential conflict of interest.

The movies for this article can be found online at: https://www.mediterranee-infection.com/acces-ressources/donnees-pour-articles/pandoravirus-massiliensis-tomograms/

