## Supplementary figures and images for "Evidence of a cellulosic layer in Pandoravirus tegument and the mystery of the genetic support of its biosynthesis"

### Supplementary Figure 1

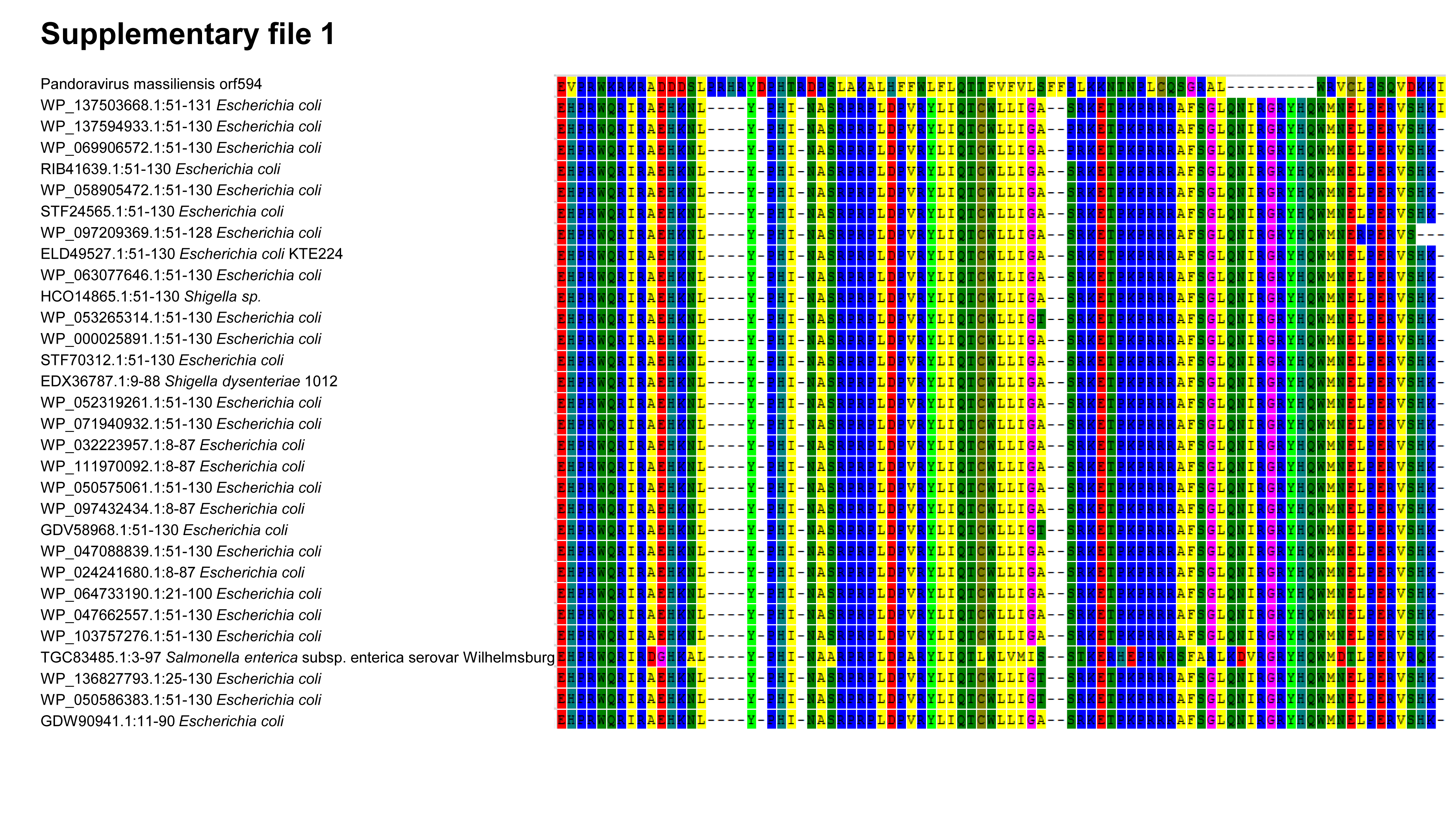
